## Supplemental Table 1 for "Identifying *in vivo* genetic dependencies of melanocyte and melanoma development"

**Table S1. Primer sequences**

| Primer name | Primer Sequence (5'-3') | Notes |
|---|---|---|
| TP53_CRISPRseq-F | CTGTGTTTGCCAGGAGTACTTG | CRISPR-seq |
| TP53_CRISPRseq-R | TATGTGTGTGTATGCGCTTTTG | CRISPR-seq |
| ptena_CRISPRseq-F | GAAGTGTTTTGAACTGCTGT | CRISPR-seq |
| ptena_CRISPRseq-R | GGAAGTCGTATTGTTACAGCT | CRISPR-seq |
| ptenb_CRISPRSeq-F | CCTTCTGAGGAATAAGCTGGAG | CRISPR-seq |
| ptenb_CRISPRSeq-R | GCAAGCTCATACCAGGTGTAAA | CRISPR-seq |
| albino_crisprseq-F | CCTGAAGGGACTGTACTTCC | CRISPR-seq |
| albino_crisprseq-R | GGGCATATTCTGTTTAAAACGACT | CRISPR-seq |
| tuba1a/c_crisprseq-F | CAACACCTTCTTCAGTGAGACC | CRISPR-seq, targets both tuba1a and tuba1c |
| tuba1a/c_crisprseq-R | ATCTTCCTTTCCTGTGATGAGC | CRISPR-seq, targets both tuba1a and tuba1c |
| tuba1a_exon1-F | CAGATTTCATAGCGTCTGACCA | validation of sgRNA |
| tuba1a_exon1-R | TGATTAAGCCCTTCAGACAGTTC | validation of sgRNA |
| tuba1a_exon2-F | GACACTTTCAACTTTTGCCCTTT | validation of sgRNA |
| tuba1a_exon2-R | GTGGAACAGCTGACGGTATGT | validation of sgRNA |
| tuba1c_exon1-F | ATAGAATTTTCCTCCTGCCTCC | validation of sgRNA |
| tuba1c_exon1-R | GCTGAATGACAAAGACAAGCAG | validation of sgRNA |
| tuba1c_exon2-F | ACGCTTGTATTGAAGGATGCT | validation of sgRNA |
| tuba1c_exon2-R | CTGCACAGATGGGTCATGAAG | validation of sgRNA |
| MitfaPCR-F | AGAATGTGAGCTTATTGGCGTT | genotyping |
| 5'homology_amplify-R | AGGACCGGGGTTTTCTTCC | genotyping |
| U6-sgRNA-F | GCGAAGATACGGCCACGG | in fusion mitf-gfp into 394-zU6-sgRNA |
| U6-sgRNA-R | GATCTAGAGGATCATAATCAGCCATAC CAC | in fusion mitf-gfp into 394-zU6-sgRNA |
| mitf-cas9-F | ATGATCCTCTAGATCCAACTTTGTATAG AAAAGTTGTGAGTGctaaca | in fusion mitf-gfp into 394-zU6-sgRNA |
| mitf-cas9-R | TGGCCGTATCTTCGCAGATCTGATCTA | in fusion mitf-gfp into 394-zU6-sgRNA |
| Ori-F | ggaagcggaacagatttaaatggtac | Geneweld plasmid construction |
| 2A-R | tggaagaaaaccccggtcctatggcttctcca | Geneweld plasmid construction |
| Cas9-F | aaccccggtcctatggcttctccacctaagaag | Geneweld plasmid construction |
| Cas9-R | ctcctaagaagaagagaaaggtgtgaactagtaatt aagtctcagccaccgttaa | Geneweld plasmid construction |
| 5'homologycheckF | CCGAGCAGAGGTGTAAAAAG | Geneweld sequence check |
| R3'_pgtag_seq | ATGGCTCATAACACCCCTTG | Geneweld sequence check |
| pPrism_VectorF | gcaggagacgtggaagaaaacc | for in fusion cloning of gBlock into pPRISM vector |
| pPrism_VectorR | ttgcccgacattatcgcgagc | for in fusion cloning of gBlock into pPRISM vector |
| 5'Fmitfa | gcggTTTCACAGGGCCCGGCCCCAGCC AGCCCGCCGAGCACGGCATGACCCCG GGA | Geneweld homology arm oligo |

| | | |
|---|---|---|
| 5'Rmitfa | gaagTCCCGGGGTCATGCCGTGCTCGGCGGGCTGGCTGGGGCCGGGCCCTGTGAAA | Geneweld homology arm oligo |
| 3'Fmitfa | aagACCCGGAGCCAGCGCTCCCAACAGCCCTATGGCCCTTCTCACCCTCAAAAA | Geneweld homology arm oligo |
| 3'Rmitfa | cggTTTTTGAGGGTGAGAAGGGCCATAGGGCTGTTGGGAGCGCTGGCTCCGGGT | Geneweld homology arm oligo |

| **Gene** | **gRNA sequence** |
|---|---|
| NT | AACCTACGGGCTACGATACG |
| *albino* | GTTTGGGAACCGGTCTGAT |
| *sox10* | GGCCGCGCGCAGGAAACTGG |
| *ptena* | AATAAGCGGAGGTACCAGG |
| *ptenb* | AGACAGTGCCTATGTTCAG |
| *p53* | GGTGGGAGAGTGGATGGCTG |
| *mitfa* | CACGGCATGACCCCGGGACC |
| *tuba1a/c* sg1(exon 2) | CGTGATCTCACCAATGACAG |
| *tuba1a/c* sg2 (exon 2) | GGTCTACAAAGACAGCCCTA |
| *tuba1a* (exon 1) | AGCAACACTACTAGAAACAA |
| *tuba1c* (exon 1) | CACTCACCATTATTCTGGAA |

| Probe name | Sequence |
|---|---|
| cas9-i1-even-01 | AGCACCTTAAATTTCTTGCTAGGGAAAgAAgAgTCTTCCTTTACg |
| cas9-i1-even-02 | TTTGTAATCTCGCTGTTCACTCTCAATgAAgAgTCTTCCTTTACg |
| cas9-i1-even-03 | TTCTTGCTGTTCTTTTCAGGCGTGTAAgAAgAgTCTTCCTTTACg |
| cas9-i1-even-04 | AGCCAGATAGATCAGTCTCAGGTCGAAgAAgAgTCTTCCTTTACg |
| cas9-i1-even-05 | CGTTATCAAATGTGCGCTGCTTCCTATgAAgAgTCTTCCTTTACg |
| cas9-i1-even-06 | GTGGCTTGCCAGATACAGGAAGTTCTAgAAgAgTCTTCCTTTACg |
| cas9-i1-even-07 | CAAACAGCTGTTTCTGTTCGTTATCAAgAAgAgTCTTCCTTTACg |
| cas9-i1-even-08 | AACTCGCTAATCTGTTCAATGATCTAAgAAgAgTCTTCCTTTACg |
| cas9-i1-even-09 | GTCTTCGACTCCGCTGATCTCCACGATgAAgAgTCTTCCTTTACg |
| cas9-i1-even-10 | ACAGGTCAATCCTTGTTTCGTACAGAAgAAgAgTCTTCCTTTACg |
| cas9-i1-even-11 | CAGAATACTTTCTTTTGAGAAACCGAAgAAgAgTCTTCCTTTACg |
| cas9-i1-even-12 | TCGAAGTACTTGAAAGCTGCAGGTGAAgAAgAgTCTTCCTTTACg |
| cas9-i1-even-13 | CCTTAATGATCTTCAGCAGATCGTGAAgAAgAgTCTTCCTTTACg |
| cas9-i1-even-14 | GTCCTCTTTGAAGGTCAGACTGTCAAAgAAgAgTCTTCCTTTACg |
| cas9-i1-even-15 | TTTTTCGATCTTCTCGCGGTTATCTAAgAAgAgTCTTCCTTTACg |
| cas9-i1-even-16 | GCGTCCTCAGCCAGATCAAAATTGCAAgAAgAgTCTTCCTTTACg |
| cas9-i1-even-17 | TTGCGCAGATGATAGATGGTTGGGTTAgAAgAgTCTTCCTTTACg |
| cas9-i1-even-18 | CGCTCTGCTTTTGTCAGGTTATCAATAgAAgAgTCTTCCTTTACg |
| cas9-i1-even-19 | GAGAAAATCTCCTGCAGGTAACAGAAAgAAgAgTCTTCCTTTACg |
| cas9-i1-even-20 | CAGCAGGGCTCCAATCAGATTTTTCAAgAAgAgTCTTCCTTTACg |
| cas9-i1-odd-01 | gAggAgggCAgCAAACggAATTGTAGTCGTCTGTAATCACTGCCC |
| cas9-i1-odd-02 | gAggAgggCAgCAAACggATATATCTGACAGCAGGATTGCGTCGG |
| cas9-i1-odd-03 | gAggAgggCAgCAAACggAACTTCGGCAGTCTCTCCAGAACCGAA |
| cas9-i1-odd-04 | gAggAgggCAgCAAACggAATTTATCAGTAGAGTCAGCCAGTTTC |
| cas9-i1-odd-05 | gAggAgggCAgCAAACggATGCAGGTCCTCTCTATTCAGTTTCAC |
| cas9-i1-odd-06 | gAggAgggCAgCAAACggTAGTACTTAGAAGGCAGGGCCAGTTCA |
| cas9-i1-odd-07 | gAggAgggCAgCAAACggAACTGGAGATCCTTTCAGCTTCTCGTA |
| cas9-i1-odd-08 | gAggAgggCAgCAAACggAATCCAGATAATGCTTGTGCTGCTCCA |
| cas9-i1-odd-09 | gAggAgggCAgCAAACggATATCGAAGCATTCAATTTTCTTGAAG |
| cas9-i1-odd-10 | gAggAgggCAgCAAACggAACGGTAATACTCTGATGGATCAGTGT |
| cas9-i1-odd-11 | gAggAgggCAgCAAACggAATGTCTGCACCTCAGTTTTCTTGACG |
| cas9-i1-odd-12 | gAggAgggCAgCAAACggAACCCAGGTTTGTCAGAGTGAACAGAT |
| cas9-i1-odd-13 | gAggAgggCAgCAAACggAAAAGCGCCCAGGCTTGCGTTAAATCT |
| cas9-i1-odd-14 | gAggAgggCAgCAAACggAAGTGAATCAGCTGCATAAAATTGCGG |
| cas9-i1-odd-15 | gAggAgggCAgCAAACggAACAGAAAGGGGTAGAAGTCTTCCTGG |
| cas9-i1-odd-16 | gAggAgggCAgCAAACggAATTGAAGTTAGGAGTCAGGCCCAGAC |
| cas9-i1-odd-17 | gAggAgggCAgCAAACggTATTTTCGTGATAGGCGACCTCGTCCA |
| cas9-i1-odd-18 | gAggAgggCAgCAAACggTATTCCTCTGAGTGATCAGTTTGGCAT |
| cas9-i1-odd-19 | gAggAgggCAgCAAACggAACTATTCTTCCTTCTGGTATAGCGCC |
| cas9-i1-odd-20 | gAggAgggCAgCAAACggAAGATGCTGTGCCTGTCGGTGTTACCC |
| mitfa-i4-even-01 | GCTCAGCTTGGCTCCCAGTGCACTGATTCTCACCATATTCgCTTC |
| mitfa-i4-even-02 | TCTCACAGTTGAGTGTGAGAAGGGCATTCTCACCATATTCgCTTC |
| mitfa-i4-even-03 | TGGCTCTGCTGGATGTGGTACTTCGTATCTCACCATATTCgCTTC |
| mitfa-i4-even-04 | CATGAGCACGGGCTTGCATTTCAAGAATCTCACCATATTCgCTTC |

| | |
|---|---|
| mitfa-i4-even-05 | GCTCTTTCTGCAATTTCCTAATGTATATCTCACCATATTCgCTTC |
| mitfa-i4-even-06 | TGTTGTAGACCCCCGGCTCGCTGTCAATCTCACCATATTCgCTTC |
| mitfa-i4-even-07 | CATCCGGTCTTTGATAAGAGTCAAAAATCTCACCATATTCgCTTC |
| mitfa-i4-even-08 | AGTTGGCATTGCTAACAGACATTGTATTCTCACCATATTCgCTTC |
| mitfa-i4-even-09 | CACCTCATGTCTGGATCATTTGACTAATCTCACCATATTCgCTTC |
| mitfa-i4-even-10 | CATTGTTGAGGTCCAAAGTGGTAGGTATCTCACCATATTCgCTTC |
| mitfa-i4-even-11 | CGAGCCACTAACTCAGCGGAATAAAAATCTCACCATATTCgCTTC |
| mitfa-i4-even-12 | TACTGTTGGAAGGTACTGGGGATCCAATCTCACCATATTCgCTTC |
| mitfa-i4-even-13 | TGGGTACAAATTGGATGTGCAATCCAATCTCACCATATTCgCTTC |
| mitfa-i4-even-14 | ATCATGCCCGGAGATGGAGTAACAGTATCTCACCATATTCgCTTC |
| mitfa-i4-even-15 | GTTTGCGTGTTCTAGTCTCTTCTGTAATCTCACCATATTCgCTTC |
| mitfa-i4-even-16 | AGGGTGTTGTCCATAAGCATGTCCTAATCTCACCATATTCgCTTC |
| mitfa-i4-even-17 | GCTGTACATGTCCAGGAGGTTTGCTATTCTCACCATATTCgCTTC |
| mitfa-i4-even-18 | GTCCTGCATCCATGAACCCAAGAATAATCTCACCATATTCgCTTC |
| mitfa-i4-even-19 | TTGAGGGGCAGGAGTTACTGATGGATTTCTCACCATATTCgCTTC |
| mitfa-i4-even-20 | GCTCTTCATGATCTGGATGTAACACAATCTCACCATATTCgCTTC |
| mitfa-i4-odd-01 | CCTCAACCTACCTCCAACATCAGGTAGTGCTTCACCTGCTGCCTC |
| mitfa-i4-odd-02 | CCTCAACCTACCTCCAACATTAGGGCTGTTTGGAGCGCTGGCTCC |
| mitfa-i4-odd-03 | CCTCAACCTACCTCCAACTAGGGGTTTCCAGGTGGGTCTGAACCA |
| mitfa-i4-odd-04 | CCTCAACCTACCTCCAACAACCTGAATACGGAGCATGAGATGTCT |
| mitfa-i4-odd-05 | CCTCAACCTACCTCCAACTACCACCGATGCTTTCAGGATGGTGCC |
| mitfa-i4-odd-06 | CCTCAACCTACCTCCAACAACCGTGGGACTGTCATTGTAGCTGAT |
| mitfa-i4-odd-07 | CCTCAACCTACCTCCAACAATGCCACATGGACCAGCTTTGTCCAT |
| mitfa-i4-odd-08 | CCTCAACCTACCTCCAACATTGTTCGTCCATACTGCTGCTGCCGC |
| mitfa-i4-odd-09 | CCTCAACCTACCTCCAACAAGGAATTAAAGTCCCCAGCTCCTTAA |
| mitfa-i4-odd-10 | CCTCAACCTACCTCCAACTAGGGACATGTCAGGACTGGGAAGGTG |
| mitfa-i4-odd-11 | CCTCAACCTACCTCCAACAACTGGAAGAAGCTACAACGGTGAGTC |
| mitfa-i4-odd-12 | CCTCAACCTACCTCCAACAAAGGACAACAGCGGGTCGCTGCTGCC |
| mitfa-i4-odd-13 | CCTCAACCTACCTCCAACAACATCCCAGGCTCCTGTTTTATTGCT |
| mitfa-i4-odd-14 | CCTCAACCTACCTCCAACTAAATTCCCTTTTGACGGCCGGCAGGT |
| mitfa-i4-odd-15 | CCTCAACCTACCTCCAACAAGTTTTCCAGCTCTTTTGCTTTCTGC |
| mitfa-i4-odd-16 | CCTCAACCTACCTCCAACAAAGCTTGGTGGAGGCCTTGTTTGGGC |
| mitfa-i4-odd-17 | CCTCAACCTACCTCCAACATAACTGGAATCGTGTTTGTCATTTGA |
| mitfa-i4-odd-18 | CCTCAACCTACCTCCAACAACATCACTGTAGCTTGATTCCAAACT |
| mitfa-i4-odd-19 | CCTCAACCTACCTCCAACTTCTCCAGCTGGAGGAAGAGCATGATT |
| mitfa-i4-odd-20 | CCTCAACCTACCTCCAACAAGTACTTACAACACACACACACACAA |

5' homology mitfa-Cas9 gblock sequence
gtagcgttgccaatgatgttacagatgagatggtcagactaaactggctgacggaatttatgcctcttccgaccatcaagcattttat
ccgtactcctgatgatgcatggttactcaccactgcgatccccggaaaaacagcattccaggtattagaagaatatcctgattcag
gtgaaaatattgttgatgcgctggcagtgttcctgcgccggttgcattcgattcctgtttgtaattgtccttttaacagcgatcgcgtatttc
gtctcgctcaggcgcaatcacgaatgaataacggtttggttgatgcgagtgattttgatgacgagcgtaatggctggcctgttgaac
aagtctggaaagaaatgcataaacttttgccattctcaccggattcagtcgtcactcatggtgatttctcacttgataaccttatttttga
cgaggggaaattaataggttgtattgatgttggacgagtcggaatcgcagaccgataccaggatcttgccatcctatggaactgc
ctcggtgagttttctccttcattacagaaacggctttttcaaaaatatggtattgataatcctgatatgaataaattgcagtttcatttgat
gctcgatgagtttttctaatcagaattggttaattggttgtaacactggcaaaaaggatctaggtgaagatcctttttgataatctcatg
accaaaatcccttaacgtgagttttcgttccactgagcgtcagaccccgtagaaaagatcaaaggatcttcttgagatcctttttttctg
cgcgtaatctgctgcttgcaaacaaaaaaaccaccgctaccagcggtggtttgtttgccggatcaagagctaccaactctttttccg
aaggtaactggcttcagcagagcgcagataccaaatactgttcttctagtgtagccgtagttaggccaccacttcaagaactctgt
agcaccgcctacatacctcgctctgctaatcctgttaccagtggctgctgccagtggcgataagtcgtgtcttaccgggttggactca
agacgatagttaccggataaggcgcagcggtcgggctgaacggggggttcgtgcacacagcccagcttggagcgaacgacc
tacaccgaactgagatacctacagcgtgagctatgagaaagcgccacggttcccgaagggagaaaggcggacaggtatccg
gtaagcggcagggtcggaacaggagagcgcacgagggagcttccagggggaaacgcctggtatctttatagtcctgtcgggtt
tcgccacctctgacttgagcgtcgatttttgtgatgctcgtcaggggggcggagcctatggaaaaacgccagcaacgcggcctttt
acggttcctggccttttgctggccttttgctcacatgttctttcctgcgttatcccctgattctgtggataaccgtattaccgcctttgagtga
gctgataccgctcgccgcagccgaacgaccgagcgcagcgagtcagtgagcgaggaagcggaacagatttaaatggtaccg
agcagaggtgtaaaaagtactcaaaaattttactcaagtgaaagtacaagtacttagggaaaattttactcaattaaaagtaaa
agtatctggctagaatcttacttgagtaaaagtaaaaaagtactccattaaaattgtacttgagtattaaggaagtaaaagtaaaa
gcaagaaagaaaactagagtggtctccctttagtgagggtcaattgatGGGAGGCGTTCGGGCCACAGCGGTTT
CACAGGGCCCGGCCCCAGCCAGCCCGCCGAGCACGGCATGACCCCGGGActtcccggagccac
gaacttctctctgttaaagcaagcaggagacgtggaag
